# Development and pharmacological evaluation of an intranasal liposomal norbinaltorphimine formulation for the prevention of pain-induced negative affect

**DOI:** 10.64898/2026.08.26.747378

**Authors:** Jesús D. Lorente, Yolanda Campos-Jurado, Miquel Martínez-Navarrete, Javier Cuitavi, Marta Cervera-Sospedra, Jessica A. Higginbotham, Ana Melero, Ana Polache, Antonio J. Guillot, Jose A. Morón, Lucía Hipólito

## Abstract

Chronic pain is frequently accompanied by negative affect and motivational deficits due to dysregulated mesocorticolimbic dopamine and kappa opioid receptor (KOR) signalling. Although intracranial KOR antagonism prevents pain-induced negative affect in preclinical models, systemic KOR antagonists can produce adverse off-target effects in the periphery, thereby limiting its clinical utility. Consistent with this, we found that systemic administration of KOR antagonist, norbinaltorphimine (NorBNI), exacerbated motivational deficits in rats with persistent inflammatory pain. We hypothesized that maximizing central and minimizing peripheral KOR antagonism could overcome these limitations. To test this, we engineered an intranasal liposomal NorBNI formulation incorporated into an in situ–forming mucoadhesive hydrogel to enable selective nose-to-brain delivery (NorBNI-LV-HG). We characterized its physicochemical properties and functional efficacy in rats with inflammatory pain produced by Complete Freund’s Adjuvant (CFA). NorBNI-loaded liposomes exhibited high drug entrapment efficiency, nanometric size, and suitable surface charge for intranasal administration. The selected thermosensitive hydrogel demonstrated appropriate gelation properties and sustained drug release. Intranasal administration of NorBNI-LV-HG produced negligible systemic NorBNI levels compared with intraperitoneal delivery. In vivo microdialysis showed that NorBNI-LV-HG prevented KOR agonist–induced reductions in nucleus accumbens (NAc) dopamine release, confirming functional central KOR blockade. Behaviourally, intranasal NorBNI-LV-HG attenuated pain-induced impairments in sucrose motivation. Importantly, unlike systemic NorBNI, repeated intranasal NorBNI-LV-HG did not alter mechanical nociceptive thresholds in pain-naïve animals, suggesting this strategy mitigates unwanted peripheral nociceptive effects. Together, these findings demonstrate that intranasal NorBNI-LV-HG achieves functional brain KOR antagonism while minimizing systemic exposure and off-target effects. Selective nose-to-brain delivery of KOR antagonists therefore represents a promising therapeutic strategy to prevent and potentially reverse the affective and motivational consequences of pain and may overcome key translational barriers associated with systemic KOR treatments.

## 1. Introduction

Chronic pain affects approximately 30% of the European population (Rometsch et al., 2025) and represents a major public health challenge because of its profound impact on the quality of life and socioeconomic burden (Bushnell et al., 2013; Ashrafioun et al., 2020). Beyond persistent nociception, pain is frequently accompanied by negative affect and motivational disturbances, including anxiety, anhedonia and anergy, which substantially impair daily functioning and contribute to poor clinical outcomes. These behavioural alterations are increasingly recognized as intrinsic components of chronic pain syndromes, like fibromyalgia (Wolfe et al., 2010). Pain-induced negative affect is linked with dysfunction of the mesocorticolimbic system (MCLS), which may increase vulnerability to substance use disorders, thereby further complicating opioid-based pain management (Bair et al., 2003; Wood et al., 2007; Dantzer et al., 2008).

The MCLS regulates reward processing, motivated behaviour and goal-directed behaviour (Everitt and Robbins, 2005; Salamone and Correa, 2012). Neuroimaging studies demonstrate that patients with chronic pain exhibit altered nucleus accumbens (NAc) activity and disrupted mesolimbic connectivity (Loggia et al., 2014; Martikainen et al., 2015), supporting dysfunction of reward circuitry as a neural substrate for pain-associated motivational deficits. Consistent with these clinical observations, preclinical studies have shown that persistent inflammatory pain suppresses dopamine (DA) neuron activity through alterations in both mu (MORs) and kappa opioid receptor (KORs) function within the ventral tegmental area (VTA) and NAc, respectively (Ozaki et al., 2002; Schwartz et al., 2014; Hipolito et al., 2015; Campos-Jurado et al., 2020; Cuitavi et al., 2023). Among these, augmented KOR signalling has emerged as a key regulator of pain-induced negative affect and motivational dysfunction (Liu et al., 2019; Massaly et al., 2019; Lorente et al., 2022a).

Interestingly, the behavioural consequences of persistent pain exhibit marked sex differences. Although inflammatory pain reduces motivated behaviour both sexes, persistent anxiety-like behaviour, anhedonia, and alcohol relapse behaviour were more pronounced in female rats (Hipolito et al., 2015; Massaly et al., 2019; Cuitavi et al., 2021; Lorente et al., 2022b), whereas pain promotes escalation of opioid self-administration in males (Hipolito et al., 2015; Higginbotham et al., 2025). Interestingly, intra-NAc administration of the selective KOR antagonist norbinaltorphimine (NorBNI) prevents part of these behavioural alterations across sexes (Massaly et al., 2019; Lorente et al., 2022b, 2024), indicating that KOR signalling represents a shared neurobiological mechanism underlying distinct pain-associated behavioural phenotypes regardless of the sex variable. These findings identify KORs as a potential target for treating the affective and motivational comorbidities of pain in both sexes.

Despite this promising preclinical evidence, the clinical translation of KOR antagonism remains challenging. Although the selective KOR antagonist aticaprant has revealed some efficacy in Phase II clinical trials for mood disorders, treatment has been associated with some adverse effects, including cardiac events and pruritus in substance-dependent patients and others (Chavkin and Koob, 2016; Jacobson et al., 2020; Hampsey et al., 2024). Moreover, systemic KOR antagonism in patients with chronic pain could potentially interfere with endogenous spinal opioid mechanisms that contribute to pain modulation, potentially limiting its therapeutic utility (Basbaum and Fields, 1984; Naser and Kuner, 2018). To overcome these barriers, novel strategies that selectively enhance central drug exposure while minimizing peripheral receptor blockade may improve efficacy and significantly decrease off-site adverse effects (Butlen-Ducuing et al., 2016; Guha et al., 2023).

Intranasal administration provides a non-invasive approach for direct drug delivery to the central nervous system, bypassing the blood-brain barrier (BBB) through the olfactory and trigeminal neural pathways (Crowe et al., 2018; Erdő et al., 2018; Giunchedi et al., 2020). Compared to systemic administration, this route minimizes peripheral drug exposure, lowers the dose required to achieve pharmacologically effective brain concentrations, provides a rapid onset of action, and improves patient acceptability (Giunchedi et al., 2020). This administration route especially benefits from the use of nano-sized entities for several reasons. First, nanoparticles allow either the protection of active ingredients or the absorption through biological barriers of drugs with unfavourable physicochemical properties. Second, they can be used to formulate hydrophobic drugs, which cannot be dissolved in aqueous-based formulations required due to biocompatibility reasons. Third, they show sustained cargo release, which leads to different advantages such as drug administration spacing or side-effects reduction, etc. Finally, nano-sized entities can be incorporated into supracolloidal systems, giving place to advance drug delivery systems that can exert multiple functionalities to improve the drug vectorization towards the pharmacological target. In this context, liposomal systems stand out among other nanosystems because of their high biocompatibility, easy preparation and scaling-up possibilities.

Here, we tested the hypothesis that systemic NorBNI administration is insufficient to prevent pain-induced motivational deficits because of peripheral actions that limit its therapeutic window, whereas selective nose-to-brain administration would enhance central KOR antagonism while minimizing peripheral exposure. To test this hypothesis, we first assessed the efficacy of systemic NorBNI in a rat model of persistent inflammatory pain. We then developed and characterized an advanced liposomal NorBNI formulation incorporated into a thermosensitive mucoadhesive hydrogel for intranasal administration, and evaluated its physicochemical properties, target engagement and behavioural efficacy as a novel strategy to selectively modulate mesolimbic KOR signalling and prevent the affective and motivational consequences of pain.

## 2. Materials and Methods

### 2.1. In vivo experiments

#### 2.1.1 Animals and chronic inflammatory pain model

A total of 122 rats, 76 Sprague Dawley rats, 8/10 weeks old (Envigo®, Barcelona, Spain), and 46 Long-Evans rats, 8/10-week-old (Envigo®, Indianapolis, USA) were used. Animals were kept in light/dark (12/12 h, light on at 6:00 a.m.) controlled cycles, temperature 23 ± 1°C, and 60% humidity. Each animal was individually housed in a standard plastic cage (42 × 27 × 18 cm^3^) with food and tap water provided *ad libitum* throughout the experimental period. Rats used in the sucrose self-administration experiment were food restricted during the experiment, see 2.1.3. Rats were housed in the animal facilities of the University of Valencia (Sprague Dawley rats), and in the animal facilities of Washington University in St. Louis (Long-Evans rats).

We selected the Complete Freund’s Adjuvant (CFA, Calbiochem) model of inflammatory pain. This model of inflammatory pain has been broadly used as a rodent model of rheumatoid arthritis that reproduces human aspects of the disease (Fischer et al., 2017). Rats were injected subcutaneously (s.c.) with CFA in one of the hind paws, as previously described (Massaly et al., 2019; Cuitavi et al., 2021; Lorente et al., 2024) after obtaining a basal measure of motivated behaviour for each animal (see below).

Animal protocols followed in this work were approved by the Animal Care Committee of the University of Valencia and were strictly adhered to in compliance with the EEC Council Directive 63/2010 and Spanish laws (RD 53/2013) and/or the Institutional Animal Care and Use Committee of Washington University in St. Louis and were in accordance with the National Institute of Health guidelines.

#### 2.1.2. Sucrose self-administration and pain model

A total of 14 male Sprague-Dawley rats and 46 Long-Evans rats (27 males and 19 females) were used to evaluate whether intraperitoneal NorBNI or intranasal novel formulation of NorBNI (NorBNI incorporated in the liposomes dispersed in the thermosensitive hydrogel, NorBNI-LV-HG) administrations, respectively, can prevent pain-induced reductions in motivated behaviour via dynorphinergic pathway modulation. We employed an inflammatory pain model (CFA) in combination with a sucrose self-administration paradigm as previously described in male SD rats (Hipolito et al., 2015) and in male and female rodents (Massaly et al., 2019).

We used an established protocol for operant sucrose self-administration under a progressive ratio (PR) schedule to assess motivation after hind paw injections of CFA or saline, as described by Hipólito et al. (2015) and Massaly et al. (2019). The experiment was conducted in operant-conditioning chambers (Med Associates) equipped with two levers positioned on the same wall, 5 cm above the floor and 12.5 cm apart. A cue light was placed 2 cm above the active lever, which was counterbalanced across animals. A food receptacle connected to a pellet dispenser was located between the levers. At the start of each session, the cue light above the active lever was illuminated. Presses on the active lever resulted in the delivery of one sucrose pellet followed by a 20-second timeout, during which the cue light was turned off and lever presses were retracted. Responses on the inactive lever were recorded but had no outcome. The protocol consisted of a training period under fixed ratio (FR) schedules, followed by a testing period using a PR schedule to assess motivation. During training, rats began on an FR1 schedule and progressed to FR2 and FR5 after meeting acquisition criteria (≥50 pellets/session for 3 consecutive sessions). FR2 and FR5 were maintained for three sessions each. Following training, a baseline PR session was conducted. The PR schedule followed the equation: ratio = [5 × e^(0.2 × infusion number)] – 5, rounded to the nearest integer (Roberts and Bennett, 1993), yielding the following response requirements: 1, 2, 6, 9, 12, 15, 20, 25, 32, 40, 50, 62, 77, 95, etc. Each session lasted 2 hours. After the baseline PR session, all animals received a CFA injection, as previously described. Following the CFA injections, two protocols were performed. The first protocol consisted of administering NorBNI (10 mg/kg) (Wille-Bille et al., 2017) or vehicle intraperitoneally just after CFA injection. Animals underwent a second PR session to assess post-treatment motivation 48 h after CFA injection.

In the second protocol, rats received once-daily intranasal administration of NorBNI-LV-HG (33 or 130 µg/kg) or vehicle (the same liposomal formulation without NorBNI, LV-HG) for four consecutive days following CFA injection. Intranasal administration was performed under isoflurane anaesthesia by pipetting no more than 50 μL of liposome-containing gel into each nostril. 24 hours after the final administration, animals underwent a second PR session to assess post-treatment motivation.

#### 2.1.3 In vivo systemic distribution assays

A total of 28 Sprague–Dawley rats (14 males and 14 females) were included in this study and were allocated to two experimental groups based on the route of NorBNI administration. One group received intranasal NorBNI-LV-HG once daily for four consecutive days at doses of 100, 130, or 200 μg/kg (n = 3–4 animals/sex/dose). As indicated above, intranasal administration was performed under isoflurane anaesthesia by carefully pipetting no more than 50 μL of the liposome-containing hydrogel into each nostril.

The reference group received NorBNI as an intravenous bolus via the lateral tail vein once daily for four consecutive days. To provide a systemic administration benchmark for comparison with the nose-to-brain formulation, animals received 2 mg/kg NorBNI, corresponding to a ten-fold higher dose than the highest intranasal dose (n = 6, 3 males and 3 females). This design enabled the evaluation of whether the intranasal formulation minimized peripheral tissue distribution compared with systemic intravenous administration.

Animals from both groups were euthanized 2 hours after the final administration by pentobarbital overdose (65 mg/kg). Following euthanasia, plasma, liver, lungs, and spleens were collected and snap-frozen in dry ice. Tissues were homogenized using a stabilising solution (44.39 g/L diammonium hydrogen citrate and 100 g/L metaphosphoric acid in dH_2_O) at a ratio of 1.5 mL stabilising solution per 0.75 mL plasma or 1 g tissue, with a Turrax homogenizer. Then the samples were centrifuged at 850G for 10 min at 4°C, and the supernatant was collected and filtered with a 0.22 µm filter. Finally, 70 μL of each sample was injected into the high-performance liquid chromatograph (HPLC) system for the detection of NorBNI in plasma and peripheral tissues according to the analytical method described in Section 2.2.3.

#### 2.1.4 Dopamine microdialysis

Microdialysis experiments were carried out in 18 rats (9 males and 9 female Sprague-Dawley rats). All surgeries were performed under isoflurane (1.5 minimum alveolar concentration, MAC) anaesthesia and under aseptic conditions. Rats were stereotaxically implanted (Stoelting, Kiel, WI) with bilateral vertical concentric-style microdialysis probes into the NAc core (NAcC; Males - anteroposterior: 1.4 mm, mediolateral: 1.5 mm and dorsoventral 7.8 mm from bregma; Females - anteroposterior: 1.2 mm, mediolateral: 1.4 mm and dorsoventral 7.6 mm from bregma) (Paxinos and Watson, 2009). These probes were constructed according to Santiago and Westerink (Santiago and Westerink, 1990), containing 2 mm of active membrane (Hospal AN69; molecular cutoff 60,000 Da).

Animals were administered intranasally either with NorBNI-LV-HG (n = 4-5/sex) or blank liposome hydrogel (n = 4/sex) as a control once daily for 4 consecutive days (starting the day of the surgery) at a dose of 100 μg/Kg. Intranasal administration was performed under isoflurane anaesthesia by pipetting no more than 50 μL of liposome-containing gel into each nostril.

Microdialysis experiments were performed as previously described (Campos-Jurado et al., 2020; Cuitavi et al., 2025). One hour after the last intranasal administration, animals were placed in Plexiglas bowls. A PE10 inlet tubing was attached to a 2.5-mL syringe (Hamilton, Spain) mounted on a syringe pump (Harvard Instruments, South Natick, MA) and connected to the probes. Probes were continuously perfused with artificial cerebrospinal fluid (aCSF) comprising 0.1 mM aqueous phosphate buffer containing 147 mM NaCl, 3.0 mM KCl, 1.3 mM CaCl_2_, and 1.0 mM MgCl_2_ (pH 7.4) at a flow rate of 3.5 mL/minute. After a minimum stabilization period of 1 h, samples were collected every 20 min, and extracellular DA levels were determined immediately after collection by using offline HPLC with electrochemical detection. The HPLC system consisted of a Waters 510 series pump in conjunction with an electrochemical detector (Mod. Decade, Antec, Leyden, the Netherlands). The applied potential was + 0.55 V (ISAAC cell; Antec, Leyden, the Netherlands). Dialysates were injected into a 2.1 mm RP-18 column (Phenomenex, Spain, Gemini-NX 3 µm, 100 x 2.00 mm) with a 65 µL sample loop. The mobile phase consisted of a sodium acetate/acetic acid buffer (0.05 mol/L, pH=6) containing 140 mmol/L of sodium chloride, 200 mg/L of 1-octanesulfonic acid, 100 mg/L of EDTA, and 150 mL/L of methanol. The mobile phase was pumped through the column at a flow rate of 0.06 mL/minute. Chromatograms were analysed and compared with standards (1.1, 2.2, 5.5, and 11 nM) run separately on each experimental day, using the AZUR 4.2 software (Datalys, France).

Once the DA baseline level was established (defined as 3 consecutive samples with 10% variation in DA content), animals were subcutaneously administered with the KOR agonist U-50488 (0.5 mg/kg). DA levels were analysed every 20 minutes for 100 minutes and area under the curve AUC were calculated for 40 minutes blocks defined as: baseline (-40 to 0 min), post-treatment 1 (from 0 to 40 min) and post-treatment 2 (from 40 to 80 min). At the end of the protocol, rats were euthanized with a pentobarbital overdose, and the brains were removed and rapidly frozen in dry ice; 40 mm-thick coronal slices of the NAc core were obtained using a cryostat and stained with a cresyl violet protocol to verify proper probe placement. After probe placement, one male rat was discarded because of incorrect probe placement. As a result, the number of animals included was 8 males and 9 females.

#### 2.1.5. Assessment of mechanical nociception by the Von Frey test

16 SD rats (8 male and 8 female) were used to assess mechanical nociceptive thresholds using the Von Frey test in order to determine whether systemic (intravenous) or intranasal administration of NorBNI alters mechanical sensitivity. Animals were allocated such that each experimental group consisted of two males and two females. The protocol started with a 15-minute habituation period to the behavioural room and boxes. Then, we manually applied 5 filaments (Aesthesio, San José, CA) following a simplified up-down method described previously (Bonin et al., 2014). A total of five Von Frey test sessions were conducted. The first session was carried out immediately prior to the intraplantar injection of either CFA or saline into the hind paw, to establish baseline mechanical nociceptive thresholds. Subsequent sessions were performed on days 1, 2, and 4 following the CFA/saline injections. Animals received either an intravenous injection of NorBNI (2 mg/kg, dissolved in saline) via the caudal vein or an intranasal administration of NorBNI-LV-HG (200 µg/kg) once daily for four consecutive days post-injection (for details in the formulation see section 2.2). As indicated above, intranasal administration was performed under isoflurane anaesthesia by pipetting no more than 50 μL of liposome-containing gel into each nostril. Drug administration occurred 30 minutes prior to each Von Frey test session. Mechanical sensitivity was expressed in grams (g).

### 2.2. Development and characterization of the NorBNI liposome intranasal formulation

#### 2.2.1 Liposome Materials

Phospholipon 90G (P90G) was purchased from Lipoid (Steinhausen, Switzerland). Cholesterol was purchased from Scharlab (Sentmenat, Spain). Hydroxy polymethyl cellulose (HPMC), Cholesterol, Sodium dodecyl sulphate (SDS), Poloxamers P-68 and P-127, and HPLC-quality methanol (MeOH) and ethanol (EtOH) were obtained from Sigma Aldrich (St. Louis, USA). Ultrapure water (UPW) was obtained by the Milli-Q purification system with resistance > 18 MΩ cm, and TOC < 10 ppb. NorBNI dihydrochloride was obtained from Sigma Aldrich (St. Louis, USA).

#### 2.2.2. NorBNI liposomes preparation

NorBNI-loaded liposomes (NorBNI-LV) were prepared using the film-hydration method. Briefly, 0.075 g of P90G and 0.016 g of cholesterol were mixed for 30 min with 50 mL of methanol. The organic solvent was then removed by rotatory evaporation for 12 h at 250 mBar and 55°C (R-100; BUCHI Ibérica, Barcelona, Spain). The resulting thin-film layer was subsequently hydrated with 10 mL of NorBNI aqueous solution (1 mg/mL) at 55°C for 1 h. Afterwards, the vesicle dispersion was subjected to ultrasonication for 2 h at 60°C and filtered 10 times through 0.45 and 0.22 µm nylon filters. Finally, NorBNI-LV dispersion was extruded 20 times through 200 nm polycarbonate membranes (LiposoFast®-Basic Extruder; Avestin, Otawa, Canada) to reduce their size and polydisperse index (PDI).

#### 2.2.3 Analytical determination and quantification of NorBNI by high-performance liquid chromatography

NorBNI levels were determined by high-performance liquid chromatography combined with electrochemical detection (HPLC-ECD). The HPLC-ECD system consisted of a Waters 510 series pump coupled with an electrochemical detector (Mod. Decade, Antec, Leyden, the Netherlands). The applied potential was +0.55 V (ISAAC cell; Antec, Leyden, the Netherlands). The mobile phase consisted of 0.1 M citric/citrate buffer pH 3.8: Methanol (65:35), conditioned with 3 mg/L of EDTA, 100 mg/L of sodium dodecyl sulfate (SDS), and 2 mM of Potassium chloride (KCl). The mobile phase was pumped at a flow of 0.7 mL/min, and a reversed-phase C18 column (Chrospher® 100 RP-18 (5 µm), 12.5×0.4 cm) was used as a stationary phase. The analytical method was validated in terms of intraday and interday precision and specificity, linearity, and accuracy within the concentration range 0.5−500 µg/mL.

#### 2.2.4. NorBNI liposomes characterization

NorBNI-LV were characterized in terms of size, PDI and ζ-potential, entrapment efficacy (EE), phospholipid recovery and in vitro drug release.

Particle size, PDI and ζ-potential were measured using a Zetasizer Nano Series (Malvern Instruments, Malvern, UK). NorBNI-LV particle size and PDI were measured by dynamic light scattering (DLS) mode, while ζ-potential was determined by laser doppler electrophoresis (LDE) mode. All measurements were conducted at room temperature, and the reported results were obtained as an average of three independent batches (n=3).

For the EE quantification, NorBNI-LVs were disrupted using a lysis mixture consisting of a SDS solution in UPW:MeOH (10 g/L SDS, 55:45). A sample of NorBNI-LV was mixed with the lysis solution in a 1:1 proportion and vigorously stirred for 1 h. The amount of Nor-BNI was then measured by HPLC-ECD as previously described. The entrapment efficiency (EE) was calculated according to the following equation (Eq. 1):

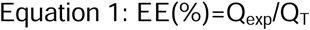

Where Q_exp_ is the final amount of NorBNI in the formulation measured by HPLC and Q_T_ is the initial amount of NorBNI used to formulate the system. The reported results were obtained as an average of three independent batches (n=3).

As a measure of the production method efficiency, the phospholipid content in liposomal dispersions was determined. For that, the Rouser et al. method was used with minor modifications (Martínez-Navarrete et al., 2024). Briefly, 100 μL of the liposomal dispersion was heated to 270 °C until solvent evaporation (OVAN SB400-E sand bath; Badalona, Spain). Afterwards, 450 μL of perchloric acid 70% (v/v) was added to the mixture and heated for 30 minutes at 250 °C. After the samples were cooled down, 3.5 mL of UPW, 500 μL of ammonium molybdate 2.5% (w/v), and 500 μL of ascorbic acid 10% (w/v) were added to the mixture, vortexed, and incubated for 7 min at 100 °C. Finally, the reaction was stopped by cooling the samples in an ice bath. The absorbance was measured by UV–Vis spectrophotometry (U-2900 spectrophotometer; Hitachi, Tokyo, Japan). The reported results were obtained as an average of three independent batches (n = 3).

#### 2.2.5. In situ hydrogel formulation

Increasing the residence time of the formulation within the nasal cavity is essential to enhance brain delivery while minimizing systemic absorption. Therefore, NorBNI-LV was incorporated into a mucoadhesive thermosensitive hydrogel designed to facilitate intranasal administration, prolong nasal retention, and provide controlled drug release. Different excipients and polymers were tested in order to achieve the best thermo-thickening and gelling properties for these purposes. NorBNI-LV-HGs were formulated by mixing different amounts of Poloxamer 127 (P-127), Poloxamer 68 (P-68), and hydroxypropyl methylcellulose (HPMC) in UPW, as described in Table 1. For the hydrogel preparation, the polymer mixture was steadily added to the NorBNI-LV dispersion under constant stirring (400 rpm) at 4°C for 10 min. Afterwards, the hydrogel was maintained at low stirring (100 rpm) overnight. Table 1 details the polymer composition of hydrogels.

**Table 1:** Polymer composition of hydrogels.

|  | P-127 (% w/v) | P-68 (% w/v) | HPMC (% w/v) |
| --- | --- | --- | --- |
| Hyg1 | 17 | - | 0.3 |
| Hyg2 | 20 | 10 | - |
| Hyg3 | 28.5 | 14.3 | - |
| Hyg4 | 15 | 2.5 | - |
| Hyg5 | 17 | 2.5 | 0.3 |

#### 2.2.6. In situ hydrogel characterization

The gelation properties of the hydrogel were addressed in terms of viscosity (Fungilab Visco Star plus viscometer). The viscosity of the different hydrogels was measured at temperatures between 5 and 50 °C. The reported results were obtained as an average of three independent batches (n=3).

#### 2.2.7. NorBNI-LV and NorBNI-LV-HG release studies

L-NorBNI-LV and NorBNI-LV-HG release studies were performed using a Franz-diffusion cell setup. For that, the receptor chamber was filled with 12 mL of UPW, and 1 mL of the formulation was added to the donor chamber. A regenerated cellulose dialysis membrane was used as a diffusional barrier between both compartments. At predefined time points, 0.25 mL samples were withdrawn from the receptor chamber, specifically at 12, 14, 16, and 18 h. In order to guarantee sink conditions, the volume of receptor media removed at every time point was immediately replaced by fresh media. The temperature remained constant at 37°C throughout the experiments. The amount of NorBNI was measured by HPLC as described before. The reported results were obtained as an average of three independent batches (n=3).

### 2.3. Statistical analysis

All the results are expressed as mean ± SEM. Before performing the statistical analysis, the Kolmogorov–Smirnov test was used to confirm a normal distribution of the data. When the groups presented a normal distribution, we used ANOVA for repeated measures followed by Bonferroni multiple comparisons for post hoc analysis. Statistical analyses were performed with IBM SPSS Statistics v24 software. The significance level was always set at p < 0.05.

## 3. Results

### 3.1. Systemic NorBNI worsens the negative affective state induced by inflammatory pain

NorBNI directly administered into the NAc shell effectively reverses negative affective states induced by pain in male and female rats. However, since the blockade of KORs might interfere with endogenous analgesia, it is possible that this route of administration does not provide the expected pharmacological effect. To solve this question, we used a well-established experimental model that reliably demonstrates pain-induced reductions in motivated behaviours driven by natural rewards (Figure 1A) (Hipolito et al., 2015; Massaly et al., 2019; Markovic et al., 2021).

**Figure 1.**
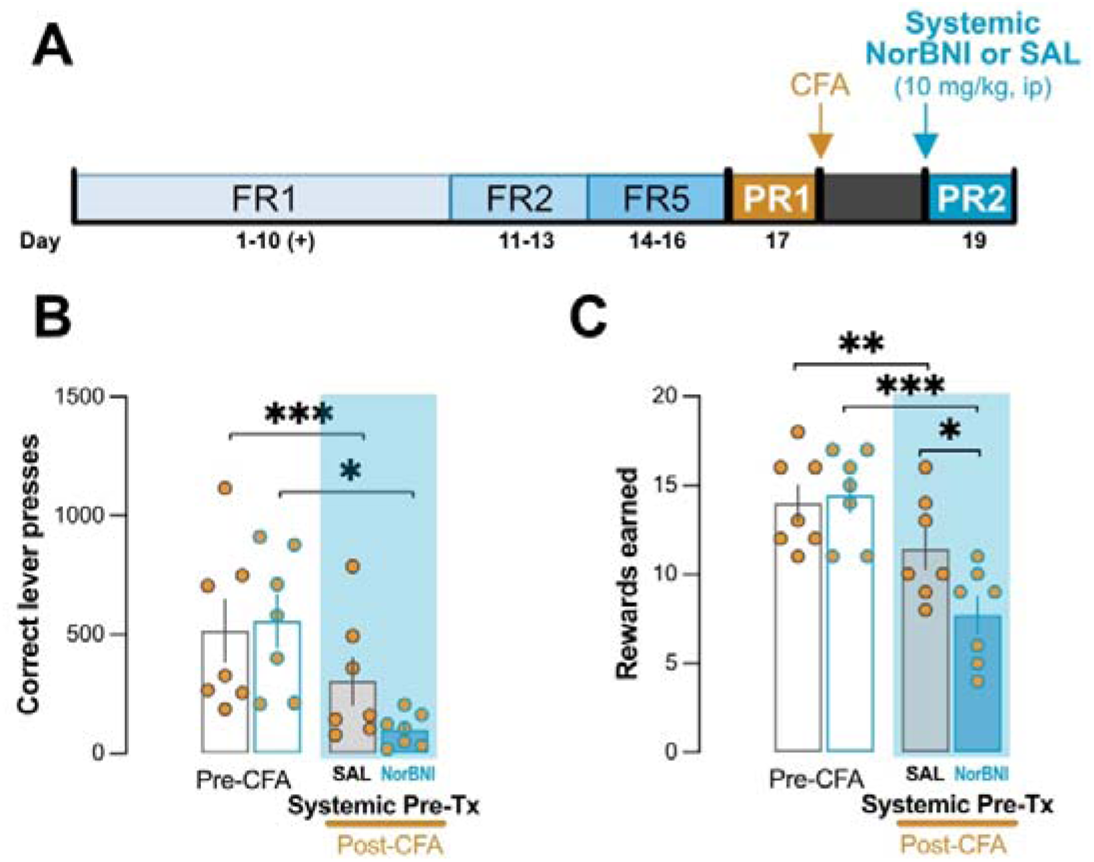
Systemic NorBNI worsens negative affect induced by pain. A) Timeline of the experimental protocol B) Number of correct lever presses performed by the control (SAL) and the NorBNI administered (nBNI 10 mg/kg) groups. C) Number of rewards earned by the SAL and the nBNI groups. Data are presented as the mean ± SEM. * denotes p < 0.05, ** denotes p < 0.01 and *** denotes p < 0.001 in the ANOVA of repeated measures followed by Bonferroni posthoc test.

Figure 1 shows that intraperitoneal administration of NorBNI produced an even greater decrease in motivated behaviour than inflammatory pain by itself. In fact, repeated measures ANOVA detected a significant interaction time*treatment variable for correct lever presses (F(1,12) = 4.547, p = 0.05) and rewards earned (F(1,12) = 13.565, p = 0.003). *Post-ho*c analyses for lever presses identified only differences between baseline and post-CFA injection in both vehicle (p < 0.05) and NorBNI (p < 0.001), without detecting any differences between groups; however, NorBNI treated animal show a non-significant decrease (p = 0.069) after treatment compared to vehicle (Figure 4B). Moreover, *post-hoc* analysis for rewards earned shows a significant reduction in both groups, vehicle (p < 0.01) and NorBNI-treated animals (p < 0.01) after the CFA injection. In addition, post-hoc analysis detected that NorBNI-treated animals decreased the rewards earned compared to vehicle (p < 0.05) after the CFA injection (Figure 4C). All these data point out that systemic NorBNI worsens the negative affective states induced by inflammatory pain. The lack of efficacy of systemic NorBNI in the presence of pain reveals the need to improve NorBNI access to the brain and reduce its presence at systemic or peripheral levels in order to assure NorBNI recovering effects on pain-induced negative affective states (Massaly et al., 2019; Lorente et al., 2024). For this reason, we developed a new formulation to administer NorBNI intranasally.

### 3.2. Development of the NorBNI liposome intranasal formulation

#### 3.2.1 NorBNI liposomes characterization

NorBNI-LV were successfully prepared using the thin-film hydration method with minimal losses during the preparation process. This was evidenced by the phospholipid recovery yield, which was 74 ± 4 % (n = 3). The entrapment efficiency was determined by measuring NorBNI levels after liposome lysis, revealing that the formulation encapsulated 90 ± 6 % of the initially formulated NorBNI (n = 3). This high loading of NorBNI in the liposomes is crucial to achieve further therapeutic levels once administered in the nasal cavity, especially due to the small volume that can be administered to the nostrils. Furthermore, the average size of NorBNI-LV was 210 ± 10 nm, the measured PDI was 0.393 ± 0.040. For its part, LDE measurements denoted that NorBNI-LV shows a ζ-potential of +4.8 ± 2.4 mV.

#### 3.2.2 *In situ-*forming hydrogel loaded with NorBNI-LV

Different formulations based on combinations of HPMC, Poloxamer 68 and Poloxamer 127 in varying proportions were used to prepare a hydrogel. Our objective was to obtain an *in-situ* hydrogel that: i) remains liquid at cold temperatures; ii) cross-links at body temperature, increasing its viscosity, and iii) presents mucoadhesive properties. All together, these properties accomplish several objectives: i) allow easy administration to the nostrils, ii) provide a long time for absorption, and iii) prevent ingestion of the formula, avoiding absorption in the gastrointestinal tract. The viscosity of the developed hydrogel prototypes was measured at temperatures ranging from 5°C to 50°C, and the results are plotted in Figure 2. Hyg1, containing only P-127 and HPMC, showed a gelation point around 25°C. Hyg2 and Hyg3, containing P-127 and P-68 in the higher concentrations, showed a rapid increase in viscosity, exceeding 10000 cPa before reaching body temperature. Hyg4 produced a slow gelation since the formulation was liquid (viscosity < 1000 cPa) above 40°C. Finally, the prototype Hyg5, maintained low viscosity (< 500 cPa) up to 35 °C, and rapidly transitioned to a gel (> 1000 cPa). Hyg1 was finally chosen based on its optimal properties for the stated objectives (this is further discussed in the discussion section).

**Figure 2.**
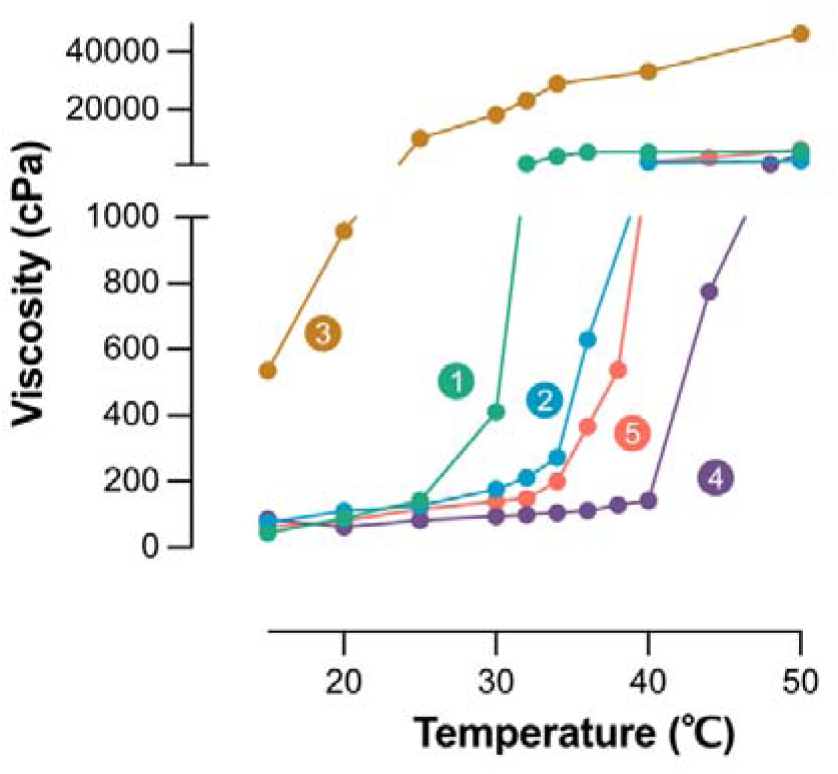
Hydrogel gelation. Viscosity measures (cPa) of the different hydrogels prototypes (1 to 5) depending on the temperature (°C).

#### 3.2.3 Assessment of NorBNI release from NorBNI-LV and NorBNI-LV-HG

To assess whether an optimal NorBNI dose is potentially available in the BioPhase, the percentage of dose delivered at 12, 14, 16, and 18 h was measured (Figure 3). A drug release of 88% of the drug at 12 h was achieved by NorBNI-LV, and this drug level remained constant onwards, denoting the full release capacity of the system. For its part, the NorBNI-LV-HG achieved a drug release percentage of around 72% after 12 h, which slightly increased in the following hours but with higher variability. Furthermore, repeated measures ANOVA were performed to compared NorBNI-LV and NorBNI-LV-HG, showing a higher release of NorBNI-LV compared to NorBNI-LV-HG (F(1,4) = 8.981, p = 0.04), however no significant differences were found in time variable (F(3,12) = 0.444, p = 0.726) and in the interaction between group*time variables were found (F(3,12) = 0.525, p = 0.584). Therefore, it can be concluded that the quantity of NorBNI release in NorBNI-LV-HG was the same but slower compared to NorBNI-LV.

**Figure 3.**
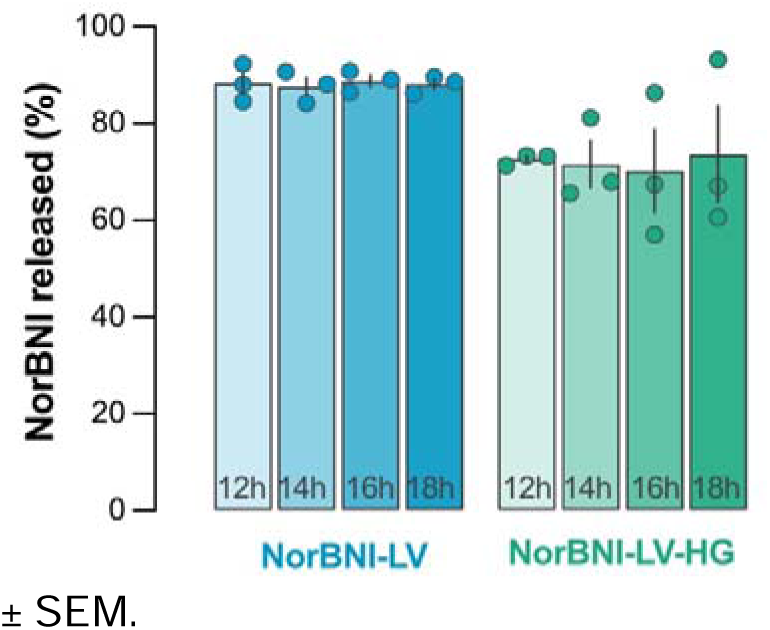
NorBNI-LV and NorBNI-LV-HG release. Representation of the NorBNI released (expressed in %) by liposomes (NorBNI-LV) and by liposomes included in the hydrogel (NorBNI-LV-HG) at different sampling times. Data are presented as the mean ± SEM.

#### 3.2.4 NorBNI *in vivo* systemic distribution

To assess whether intranasal administration of NorBNI-LV-HG reduced systemic tissue distribution relative to intravenous administration, we compared NorBNI levels in the plasma, spleen, liver, and lungs of two groups: a positive control group intravenously administered with 2 mg/kg of NorBNI per day during 4 days and three groups administered intranasally with NorBNI-LV-HG once daily for four days at three different doses (100, 130, or 200 μg/kg). Interestingly, the electrochemical HPLC method did not detect NorBNI in any tissue following intranasal administration of 100 and 130 μg/kg NorBNI, indicating that its levels were below the detection limit (74 pg/mL) and thus negligible, if present. In contrast, we were able to detect NorBNI in plasma samples of the animals administered intranasally with 200 μg/kg NorBNI NorBNI-LV-HG, although it was not posible to measure them since its levels were below the quantification limit (7.81 μg/mL). Intravenous administration of NorBNI resulted in measurable plasma levels of 17.31 ± 4.37 μg/mL. NorBNI was also detected in the liver of intravenously treated animals, although the levels were below the quantification limit.

### 3.3. Formulation efficacy

#### 3.3.1. Intranasal NorBNI-LV-HG administration blocks the kappa opioid receptor agonist effect in the NAc

Our data indicate that the effects of NorBNI on pain-induced motivational deficits depend on its access to the NAc, while peripheral actions may produce opposing effects. Accordingly, our formulation was designed to enhance NorBNI delivery to the NAc while minimizing peripheral exposure.

To assess whether the intranasal formulation achieved functional target engagement in the NAc, we performed in vivo microdialysis to measure extracellular DA levels in the NAc. DA levels were monitored before and after subcutaneous administration of the KOR agonist U-50488 (0.5 mg/kg) in rats previously treated with intranasal NorBNI (100 μg/kg) or vehicle. This approach allows direct assessment of KOR-mediated modulation of mesolimbic DA transmission at the site of action.

No sex differences were observed in the repeated-measures ANOVA for sex variable (F(1,21) = 4.042, p = 0.057), or the interaction of sex × treatment (F(1,21) = 1.048, p = 0.318), or sex × time (F(1,147) = 0.777, p = 0.608), or sex × time × treatment (F(1,147) = 0.555, p = 0.734) allowing us to pool data from males and females for subsequent analyses.

As shown in Figure 4B, extracellular DA levels changed over time following U-50488 administration. Repeated-measures ANOVA revealed significant differences in time (F(7,161) = 3.058, p = 0.002), treatment (F(1,23) = 41.005, p < 0.001), and time × treatment interaction (F(7,161) = 5.964, p < 0.001). Post hoc analyses showed that vehicle-treated animals exhibited a marked reduction in DA levels after U-50488 injection, whereas this effect was prevented in animals treated with intranasal NorBNI-LV-HG.

**Figure 4.**
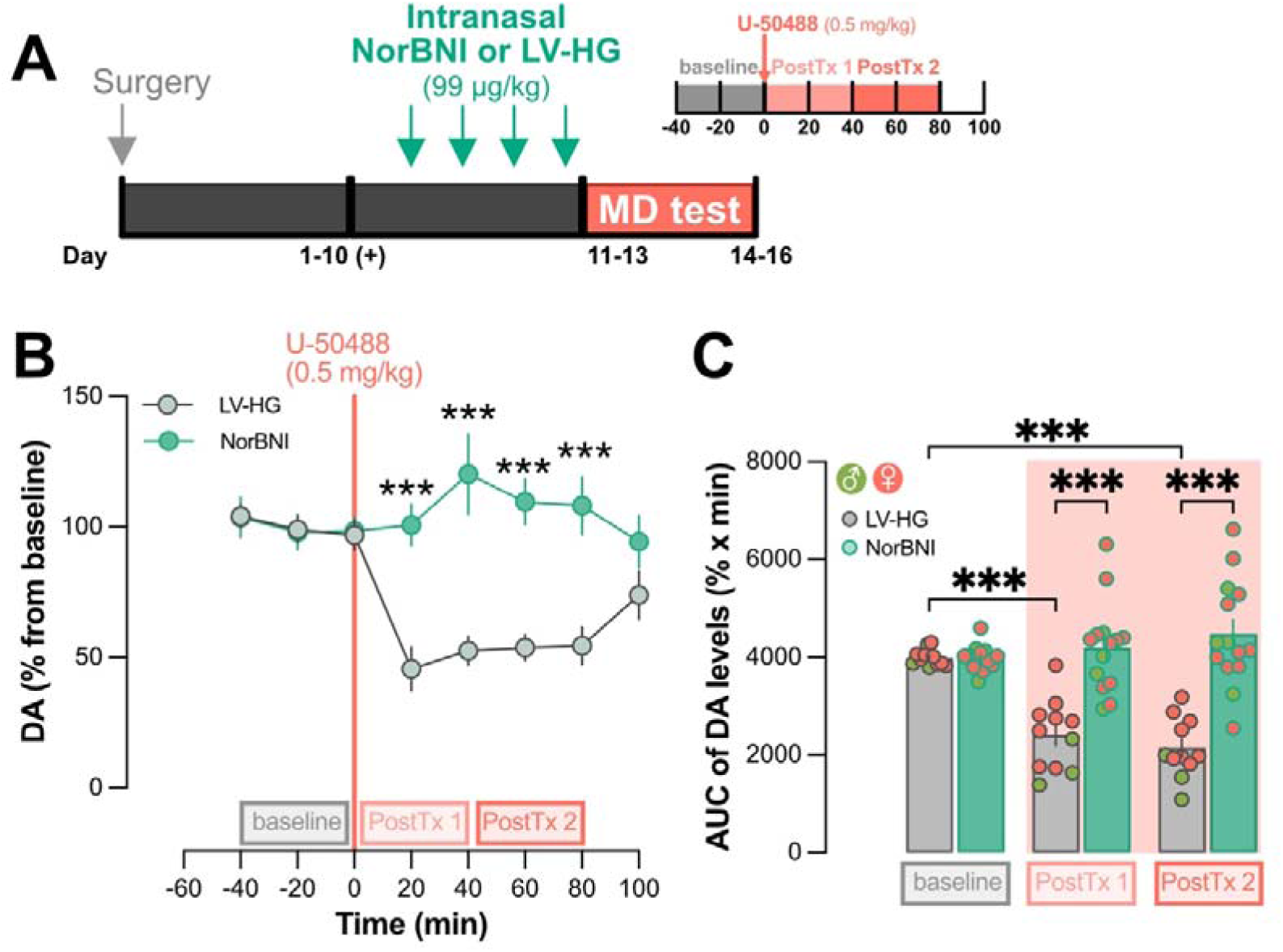
NorBNI-LV-HG prevent NAc DA reduction in NAc core induced by U-50488. A) Timeline of the protocol B) Evolution of the DA extracellular levels, represented as a percentage from baseline, in the animals treated with LV-HG (in grey) and the ones receiving NorBNI-LV-HG (NorBNI, in green) before and after the administration of 0,5 mg/kg of U-50488. C) AUC of the DA extracellular levels in the three stablished 40 min time-blocks: baseline, post-treatment 1 (PostTx1) and post-treatment 2 (PostTx 2) for the animals receiving LV-HG (grey) and NorBNI-LV-HG (green). Data points from males (green) and female (orange) rats are distinguished in 4C. Data are presented as the mean ± SEM. *** denotes p < 0.001 in the Bonferroni multiple comparisons post-hoc test.

Consistent with these findings (figure 4C), analysis of the AUC calculated in three stablished time intervals (baseline, post-treatment 1 and post-treatment 2), revealed significant differences in time (F(7,161) = 10.165, p < 0.001), treatment (F(1,23) = 40.267, p < 0.001), and treatment × time interaction (F(2,46) = 24.91, p < 0.001). Post hoc comparisons indicated that only vehicle-treated animals showed a significant reduction in DA AUC following U-50488 administration compared to pre-injection levels, whereas intranasal NorBNI-LV-HG prevented this decrease.

#### 3.3.2. Intranasal NorBNI-LV-HG administration prevents pain-induced reduction in motivation without affecting nociception

As described above, intranasal NorBNI-LV-HG effectively prevents KOR agonist-induced reduction in DA extracellular levels, indicating effective central target engagement at a 100 μg/kg dose.

To determine whether the observed effect in the BioPhase is translated into behavioural pharmacological effects, we evaluated the dose-dependent efficacy of NorBNI-LV-HG (Figure 5A) in a well-established rat model of pain-induced motivational deficits. This model combines intraplantar CFA administration with sucrose self-administration under a progressive ratio schedule of reinforcement (Schwartz et al., 2014; Hipolito et al., 2015; Massaly et al., 2019; Markovic et al., 2021). Dose selection was based on both the biodistribution and pharmacodynamic studies. Specifically, 130 μg/kg was chosen as the highest intranasal dose that showed no detectable systemic distribution, while remaining above the previously established pharmacologically effective dose of 100 μg/kg in the nucleus accumbens microdialysis assay. This strategy maximized the likelihood of detecting behavioural efficacy while minimizing peripheral exposure. In addition, a four-fold lower dose (33 μg/kg) was included to further evaluate the dose-response relationship.

**Figure 5.**
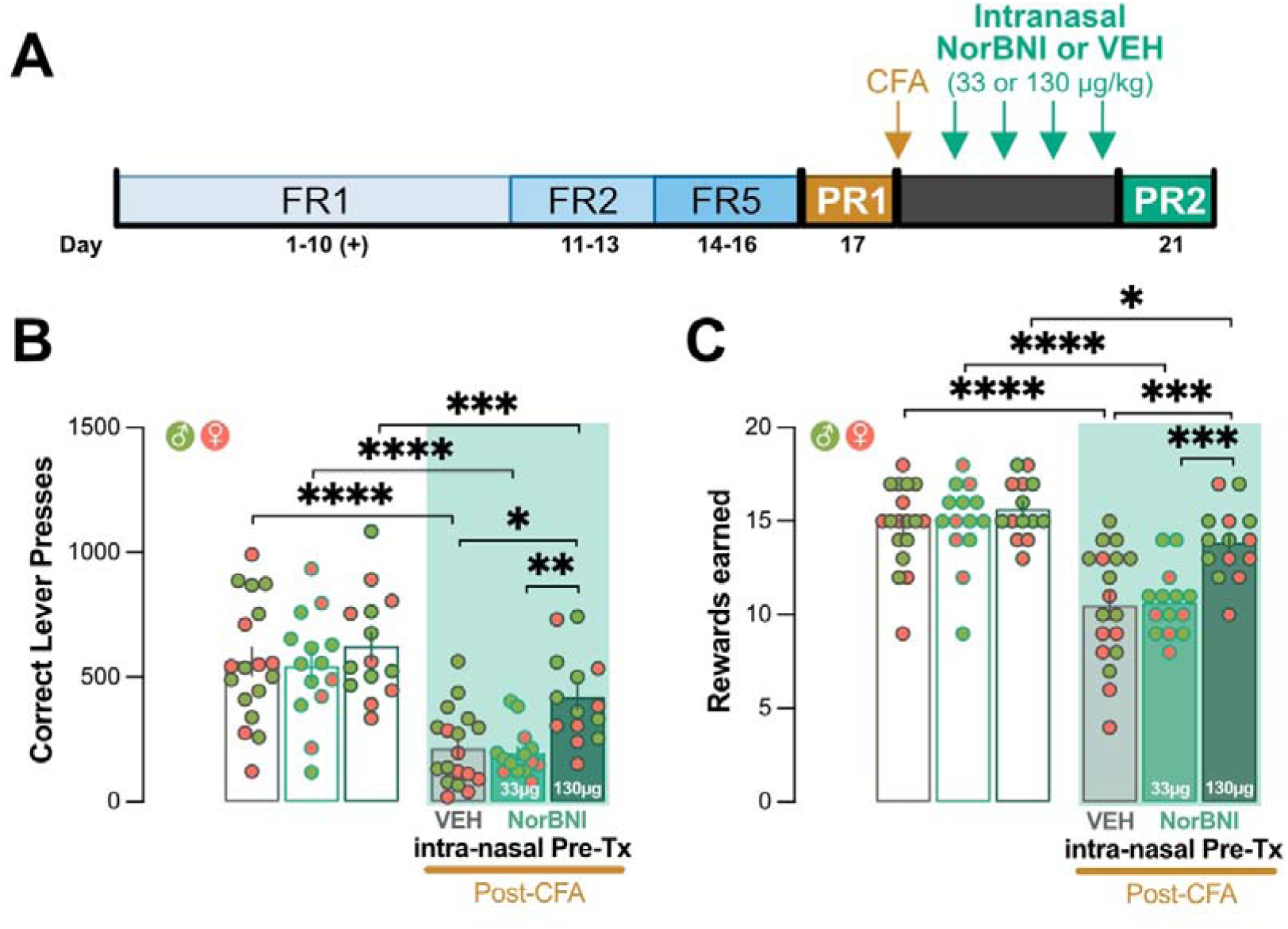
NorBNI-LV-HG dose-dependently prevent pain-induced decreased in motivation. A) Timeline of the experimental protocol B) Number of correct lever presses performed by control rats receiving LVHG (VEH) and NorBNI-LV-HG (NorBNI) administered rats at 33 or 130 μg/kg. C) Number of rewards earned by control rats receiving LVHG (VEH) and NorBNI-LV-HG (NorBNI) administered rats at 33 or 130 μg/kg. Data are presented as the mean ± SEM. Individual data points are represented in green for male rats and in orange for female rats. * denotes p < 0.05, ** denotes p < 0.01 and *** denotes p < 0.001 in the Bonferroni multiple comparisons post-hoc test.

As figure 5 shows, repeated measures ANOVA detected a significant interaction between CFA x Treatment variables for rewards earned (F(2,43) = 6.5, p = 0.0033) and a significant effect of treatment for correct lever presses (F(2,43) = 3.6, p = 0.0340) and rewards earned (F(2,43) =5.2, p =0.0093). *Post hoc* analyses indicated that, relative to baseline, animals in all groups—including the control group and both NorBNI-LV-HG treatment groups—exhibited a significant reduction in active lever presses and rewards earned. This indicates that pain-induced reduction in motivation is present in all groups. However, *post-hoc* analysis pointed out that this reduction was less pronounced in the groups that received the highest dose of NorBNI-LV-HG. In fact, animals that received the high dose of NorBNI-LV-HG, 130 μg/kg, showed statistically higher presses of the active lever and reward obtained compared to control and low dose groups, which points out a higher motivation for natural reward compared with control and low dose groups.

#### 3.3.3. Intranasal NorBNI-LV-HG administration avoids systemic NorBNI-induced alteration of nociception

Finally, to discard effects on the nociceptive threshold of our formulation, we performed a nociceptive mechanical test after repeated intranasal administration of the highest dose of NorBNI-LV-HG (200 ug/kg). Interestingly, repeated administration of a high dose in control animals does not significantly alter mechanical nociception, while the i.v. administration of NorBNI significantly reduces mechanical nociception in pain-naïve animals (Figure 6B). In fact, the ANOVA for repeated measures detected differences in the number of administration variable (F(3,18) = 6.027; p = 0.005), the type of administration variable (F(1,6) = 8.370; p = 0.028), and the interaction of both variables (F(3,18) = 6.595; p = 0.003).

**Figure 6.**
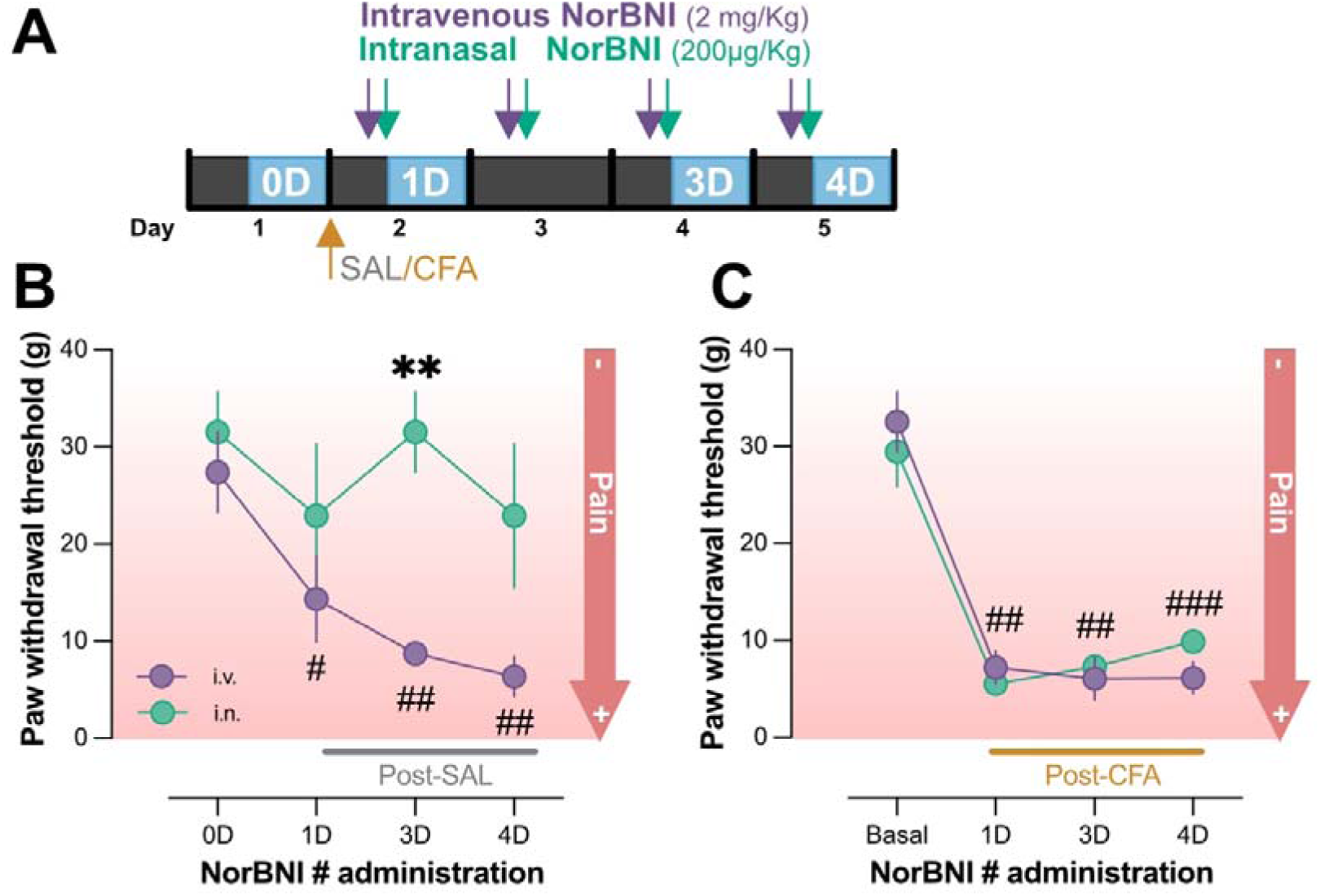
Systemic NorBNI, but not intranasal NorBNI-LV-HG (NorBNI), reduce nociception threshold in non-pain animals. A) Timeline of the behavioral experiment. B) Paw withdrawal thresholds (g) for saline-injected rats treated with systemic NorBNI (i.v.) and intranasal NorBNI-LV-HG (i.n.). ** denotes p < 0.01 i.n. vs i.p., # denotes (p < 0.05) i.v. basal vs i.v. after NorBNI administration, ## denotes (p < 0.01) i.v. basal vs i.v. after NorBNI administration (ANOVA repeated measures followed by Bonferroni multiple comparisons post-hoc test). C) Paw withdrawal thresholds (g) for CFA-injected rats treated with systemic NorBNI (i.v.) and intranasal NorBNI-LV-HG (i.n.). ## denotes (p < 0.01) pre-CFA injection vs post-CFA-Injection, ### denotes (p < 0.001) pre-CFA injection vs post-CFA-Injection (ANOVA for repeated measures followed by Bonferroni post-hoc test). Data are presented as the mean ± SEM.

As shown in Figure 6B, *post hoc* analyses revealed that pain-naïve animals receiving intravenous NorBNI exhibited a significant reduction in their mechanical nociceptive threshold beginning after the first administration and persisting throughout the protocol, relative to their baseline values. In contrast, animals treated with intranasal NorBNI-LV-HG did not show any reduction in nociceptive threshold at any point during the experimental timeline. Moreover, after three consecutive days of treatment, the NorBNI-LV-HG group displayed significantly higher nociceptive thresholds compared to the intravenously treated group. Finally, under pain conditions, we did not find any difference between the administration of intraperitoneal NorBNI and intranasal NorBNI-LV-HG (200 ug/kg). In fact, ANOVA for repeated measures did not detect differences in the type of administration variable (F(1,6) = 0.375; p = 0.563) and the interaction of both variables (F(3,18) = 1.706; p = 0.201), but it detected differences in the number of administration variable (F(3,18) = 67.107; p = 0.201). As shown in Figure 6C, *post hoc* analyses revealed that both groups together showed a significant reduction in their mechanical nociceptive threshold beginning after the first administration and persisting throughout the protocol, relative to their baseline values, without showing differences between them at any time point. This indicates that the reduction in nociceptive thresholds is associated with the administration of CFA, and no differences were found depending on the type of administration of NorBNI.

## Discussion

In this study, we demonstrate that peripheral drug exposure is a major obstacle limiting the therapeutic application of KOR antagonist for the treatment of pain-associated affective and motivational impairments. Whereas systemic NorBNI exacerbated motivational deficits and increased nociceptive sensitivity, selective nose-to-brain delivery using the developed NorBNI-LV-HG advanced intranasal formulation preserved central pharmacological activity while limiting systemic exposure. Together, our data establish selective nose-to-brain delivery as a promising strategy to overcome one of the principal translational barriers limiting the therapeutic use of KOR antagonists.

To improve the pharmacological profile of NorBNI, we selected the intranasal route because it provides a unique opportunity to reduce systemic exposure while enabling direct access to the brain through the olfactory and trigeminal pathways, thereby bypassing the BBB. In addition to this biopharmaceutical advantage, intranasal delivery is non-invasive, offers a rapid onset of action, and is particularly attractive for CNS-active compounds. However, successful nose-to-brain delivery requires pharmaceutical formulations specifically designed to maximize nasal residence time and drug availability within the limited administration volume. We therefore developed a liposomal NorBNI formulation incorporated into a mucoadhesive in situ-forming thermosensitive hydrogel (NorBNI-LV-HG) to combine efficient drug encapsulation with prolonged nasal residence time and sustained drug release.

The physicochemical properties of the formulation support its suitability for intranasal administration. High encapsulation efficiency allows therapeutically relevant amounts of NorBNI to be delivered within a small volume and improves efficiency in the production process. In addition, the nanometric particle size is compatible with efficient interaction with the nasal mucosa and has previously been associated with improved nasal permeability and nose-to-brain delivery (Sun et al., 2020). Although the liposomes displayed a moderately heterogeneous size distribution, similar PDI values have been reported for successful intranasal liposomal formulations (Youssef et al., 2018; Touitou et al., 2020; Gadhave et al., 2021). Likewise, the near-neutral ζ-potential provided by soybean phosphatidylcholine (Carreras et al., 2020) may represent an optimal balance between colloidal stability, biocompatibility and reduced epithelial toxicity compared with highly positively surface-charged systems (Formica et al., 2022; Guillot et al., 2023).

To further enhance nasal residence time, liposomes were incorporated into a thermoresponsive mucoadhesive hydrogel. The combination of HPMC and poloxamers provided appropriate gelation behaviour and viscosity for intranasal administration, allowing the formulation to remain easily applied at room temperature while rapidly gelling under physiological conditions (Cao et al., 2009; Gänger and Schindowski, 2018; Chen et al., 2019). Incorporation into the hydrogel also produced a more sustained drug release without substantially compromising the total amount of NorBNI released, consistent with the reservoir effect described for polymeric matrices (Nie et al., 2011).

Although these physicochemical properties support the suitability of the formulation for intranasal delivery, the main objective was not simply to facilitate nasal administration but also to improve the therapeutic utility of NorBNI by restricting its distribution mainly to the brain. Consistent with this objective, NorBNI was undetectable in plasma and peripheral tissues following intranasal administration at the low and intermediate doses. Even at the highest intranasal dose (200 μg/kg), NorBNI was detected only at trace levels below the limit of quantification, in marked contrast to intravenous administration, which resulted in measurable plasma and hepatic concentrations. These findings demonstrate that intranasal administration markedly limits systemic exposure to NorBNI. This reduction is particularly relevant considering the pharmacological profile of systemic NorBNI. As shown in the present study, and consistent with previous reports (Basbaum and Fields, 1984; Schepers et al., 2008; Campillo et al., 2011; Jacobson et al., 2020), systemic KOR antagonism alters nociceptive responses, an effect that may complicate its therapeutic application in chronic pain. Therefore, minimizing peripheral exposure represents an important pharmacological advantage of the formulation developed here.

Reduced systemic exposure alone does not establish therapeutic potential. Pharmacodynamic studies were therefore required to determine whether the formulation retained sufficient pharmacological activity following intranasal administration. Using in vivo microdialysis as a functional pharmacodynamic assay, we demonstrated that intranasal NorBNI (100 μg/kg) prevented the reduction in extracellular DA induced by the KOR agonist U-50488 within the NAc. Rather than simply demonstrating regional brain distribution, these findings provide direct evidence that intranasally administered NorBNI functionally antagonizes KOR signalling within a key component of the mesolimbic reward circuitry. Importantly, this effect was achieved using an intranasal dose approximately 100-fold lower than that previously required following systemic administration to produce comparable pharmacological effects (Maisonneuve et al., 1994).

The pharmacodynamic relevance of these findings is particularly evident in the context of inflammatory pain. Hyperactivity of the dynorphin/KOR system within mesolimbic circuits has been consistently associated with reduced dopaminergic neurotransmission and the development of pain-induced motivational deficits and anxiety-like behaviours (Massaly et al., 2019; Lorente et al., 2024). Previous studies have shown that direct intra-accumbal administration of NorBNI reverses these maladaptive behavioural responses (Massaly et al., 2019), establishing the importance of local KOR antagonism within this circuitry. Our findings extend these observations by demonstrating that selective intranasal delivery can achieve comparable functional KOR antagonism without the systemic exposure associated with conventional routes of administration.

Demonstrating pharmacodynamic activity alone, however, is not sufficient to support therapeutic potential. It was therefore essential to determine whether pharmacologically relevant brain KOR antagonism translated into efficacy in a relevant pathophysiological context. Using a well-established model of inflammatory pain-induced motivational deficits (Hipolito et al., 2015; Massaly et al., 2019; Markovic et al., 2021), intranasal NorBNI-LV-HG significantly dose-dependently attenuated the reduction in sucrose self-administration produced by CFA-induced inflammatory pain. These findings indicate that restoration of mesolimbic DA signalling translates into improved motivated behaviour, supporting the concept that selective central KOR antagonism directly targets one of the principal neurobiological mechanisms underlying pain-associated negative affect.

Importantly, this therapeutic efficacy was achieved without the increase in nociceptive sensitivity observed following systemic NorBNI administration. Whereas intravenous NorBNI reduced mechanical nociceptive thresholds, consistent with previous reports demonstrating that systemic KOR antagonism interferes with endogenous analgesic mechanisms (Basbaum and Fields, 1984; Schepers et al., 2008; Campillo et al., 2011; Jacobson et al., 2020), repeated intranasal administration did not alter mechanical nociceptive thresholds in pain-naïve rats. Together, these findings indicate that selective brain delivery substantially improves the therapeutic profile of NorBNI by preserving beneficial central actions while minimizing peripheral pharmacological effects.

Implications of these findings extend beyond NorBNI itself. Selective nose-to-brain delivery may represent a general strategy for improving the therapeutic utility of centrally acting KOR antagonists whose clinical development has been affected by systemic adverse effects. By increasing brain selectivity while reducing peripheral receptor blockade, this approach may permit lower therapeutic doses, improve target engagement and enhance the safety profile of KOR antagonists for disorders characterized by dysregulated dynorphin/KOR signalling, including chronic pain, depression and substance use disorders.

In conclusion, this work provides preclinical proof of concept that an appropriately designed intranasal pharmaceutical formulation offers a strategy to improve the pharmacological profile of NorBNI by enabling pharmacologically relevant brain KOR antagonism while minimizing systemic exposure and peripheral pharmacological effects associated with conventional systemic administration. These findings establish intranasal delivery as a promising strategy to prevent and reverse the affective and motivational consequences of pain and provide a framework for the future development of brain-selective KOR antagonists for neuropsychiatric disorders involved dysregulated KOR signalling.

## Acknowledgements

We would like to thank Ms. Pilar Laso and the staff from the “Servei d’Investigació-UV” for grant management. We would also thank the personnel of the Animal facilities (SCSIE) at the University of Valencia for their help and effort in assuring animal welfare.

## Author contribution

Conceptualisation: LH; Methodology: JDL, AP, JAM, AM, LH; Formal analysis: JDL, MM-N, JC, AJG, YC-J, JAH; Investigation: JDL, MM-N, JC, AJG, MC-S, YC-J, HJ ES; Writing—original draft: JDL, MM-N, JC; Resources: JAM, AM, LH; Supervision: AJG, AP, JAM, AM, LH; Writing—review and editing: JDL, MM-N, JC, AJG, AP, JAM AM, LH. All authors contributed to the article and approved the submitted version.

## Data availability statement

The raw data supporting the conclusions of this article will be made available by the authors, without undue reservation.

